# Dual-portal ligand transport in human cytochrome P450 17A1

**DOI:** 10.64898/2026.10.19.760170

**Authors:** Yingtong Xu, Adam Zalewski, Amit V. Pandey

**Author notes:** Correspondence: Prof. Dr. Amit V. Pandey. These authors contributed equally to this work (Yingtong Xu, Adam Zalewski).

## Abstract

Human cytochrome P450 17A1 (CYP17A1) is the qualitative regulator of steroidogenesis, catalyzing both 17α-hydroxylation and 17,20-lyase cleavage at a single active site, and is the target of the prostate cancer drug abiraterone. Crystal structures show an almost fully enclosed site and do not reveal how lipophilic steroids reach it from the endoplasmic reticulum membrane or how the more polar products leave. We therefore asked which routes connect the CYP17A1 active site to the protein surface, and which of them are required. Molecular dynamics of the CYP17A1 enzyme followed by CAVER tunnel clustering produced 99 transport pathways in 14 clusters, which grouped by the elements forming their mouths into four topologically distinct portals (P-I to P-IV). Five structure-guided variants of CYP17A1 were expressed in HEK293 cells and both hydroxylase and lyase reactions were measured. Occluding the dominant P-I/P-II bottleneck (A105L) left both reactions intact and slightly raised them (116.1 ± 8.0% hydroxylase, P = 0.0043; 134.8 ± 27.9% lyase, P = 0.028). A tryptophan in the F/G lumen (L242W) was tolerated (96.6 ± 6.9% and 89.9 ± 10.4%, neither significant), whereas an engineered I205C/R239C pair across the same route abolished catalysis (below 2%, P < 0.001). Both perturbations of the membrane-facing P-IV portal cut activity below 2% of wild type (P < 0.001), and CYP17A1 protein levels accounted for neither phenotype. These data support a dual-portal model of vectorial ligand transport in CYP17A1.

## 1. Introduction

Cytochrome P450 17A1 (CYP17A1, P450c17) sits at the branch point between glucocorticoid and androgen biosynthesis in the human adrenal cortex and gonads [1]. The CYP17A1 catalyzes two chemically distinct reactions at one active site: 17α-hydroxylation of pregnenolone and progesterone, and the subsequent 17,20-lyase cleavage yielding dehydroepiandrosterone (DHEA) and as a minor reactions in humans, androstenedione [1, 2]. Loss of both CYP17A1 activities causes 17α-hydroxylase/17,20-lyase deficiency [3], whereas selective loss of the lyase reaction produces isolated 17,20-lyase deficiency, reported for E305G, R347H, R358Q and V366M mutations [4–6]. Conversely, intratumoral androgen synthesis sustains castration-resistant prostate cancer, and abiraterone acetate improves survival, although CYP17A1 levels in prostate cancer cells are much lower [7–9]. Abiraterone coordinates the heme iron and inhibits both CYP17A1 reactions causing secondary glucocorticoid deficiency, so a lyase-selective inhibitor is still required [10–12].

Molecular basis of steroid substrate access remains unresolved. How does a steroid reach the buried active site of CYP17A1, and by what route does the product exits is not known. Structures of CYP17A1 with inhibitors [13, 14], and substrates [15] show a narrow, almost fully enclosed site in closed, relaxed states in which the transient openings used for ligand passage are invisible [16]. Microsomal P450s are anchored to the endoplasmic reticulum by an N-terminal helix and the catalytic domain is partly inserted into the lipid bilayer through the F/G region and the adjacent β-sheet domain [17, 18]. Membrane bound NADPH P450 oxidoreductase is required for CYP17A1 activities, and in addition, the 17,20 lyase activity also requires cytochrome b5 [19–24]. Because pregnenolone and progesterone are in membranes but the 17-hydroxylated intermediates and DHEA release into the cytosol, we investigated if a vectorial transport exists in CYP17A1 with distinct entry and exit routes rather than one bidirectional channel. The P450 tunnel literature indicates pathway openings depend on isoform, ligand and membrane, and entry and exit routes need not be open together [25, 26]. Here we mapped the tunnel ensemble of the CYP17A1 enzyme, grouped the clusters into topologically coherent portals, and probed each portal by steric and covalent perturbation.

## 2. Materials and methods

Detailed procedures, reagents and parameters are in the Supplementary Methods (Supplementary Materials).

### 2.1. Simulation, tunnel detection and portal assignment

The catalytic domain was taken from the CYP17A1-abiraterone complex (PDB 3RUK, chain D) [13], cross-referenced to the substrate complexes 4NKX and 4NKZ [15]. Residues preceding Arg45 were removed and one ordered water adjacent to the heme retained. The apo enzyme was simulated with AMBER at 310 K with explicit solvent under YASARA [27, 28]; and 83 snapshots were used in analysis. Tunnels were computed with CAVER 3.0 with heme iron as starting point [29]. Clusters whose openings were bounded by the same secondary-structure elements were grouped into portals, radius profiles were interpolated onto a common normalized coordinate, and the five standardized descriptors were analyzed by principal components (Supplementary Tables S1-S3, Supplementary Figure S1). Lining occupancy, bottleneck recurrence, core-lining composition, residue annotation and variant design are in Supplementary Tables S4-S8. Cysteine pairs were evaluated by Cβ-Cβ distance and by disulfide prediction [30].

### 2.2. Expression and activity assays

Variants were generated by site-directed mutagenesis and verified by Sanger sequencing. HEK293 cells were transfected transiently and harvested 48 h later as described [31]. 17α-Hydroxylase activity was measured by conversion of progesterone to 17α-hydroxyprogesterone and 17,20-lyase activity and conversion of 17α-hydroxypregnenolone to DHEA as previously described [32–34]. Each experiment included technical replicates and wild-type, empty-vector and untransfected controls, and activities are calculated as percentage of the wild-type control assayed in parallel in the same experiment (Supplementary Table S9, Supplementary Figure S2). Variants retaining measurable activity were carried through further rounds of assay, which is why more data points are available for them. CYP17A1 was quantified by western blot against β-actin in the same experiments.

### 2.3. Statistics

Variant-level data are the mean ± SD of independent experiments. Each variant was compared with the wild-type reference, defined as 100% in every experiment, by a two-sided one-sample t-test: ns (P ≥ 0.05), * (P < 0.05), ** (P < 0.01), *** (P < 0.001). Expression-corrected activity is measured activity divided by the immunoblot ratio of the matching lysate, × 100. Analyses ran in Python 3.12 (Supplementary Tables S10-S12).

## 3. Results and discussion

### 3.1. The CYP17A1 tunnel ensemble resolves into four constricted portals

Across 83 MD snapshots CAVER provided 99 transport pathways in 14 clusters. Grouping them by the elements forming their entry points gave four topologically distinct portals and one singleton (Figure 1, Supplementary Tables S1 and S2). P-I (clusters 1, 3, 6; 46 tunnels, 46.5%) exits between the B’-C loop and the G helix; P-II (clusters 2, 4; 35 tunnels, 35.4%) opens on the B-C loop surface; P-III (clusters 5, 8, 9, 12, 13; 11 tunnels, 11.1%) is the long, curved tunnel between the F and G helices; and P-IV (clusters 7, 10, 14; 6 tunnels, 6.1%) runs parallel to the heme plane, past Val366, into the β-domain and the membrane-facing surface. In the established nomenclature [25] these correspond to the 2b, 2e, 2a and 4/5 families. P-I and P-II dominate, open in 54.2% and 42.2% of snapshots against 13.3% and 7.2% for P-III and P-IV. Principal component analysis separate the portals along one dominant axis (PC1, 65.1%; PC2, 21.0%), contrasting short, wide, high-throughput tunnels with long, curved, narrow ones; as the mutagenesis shows, this geometric distinction does not coincide with the functional one.

**Figure 1.**
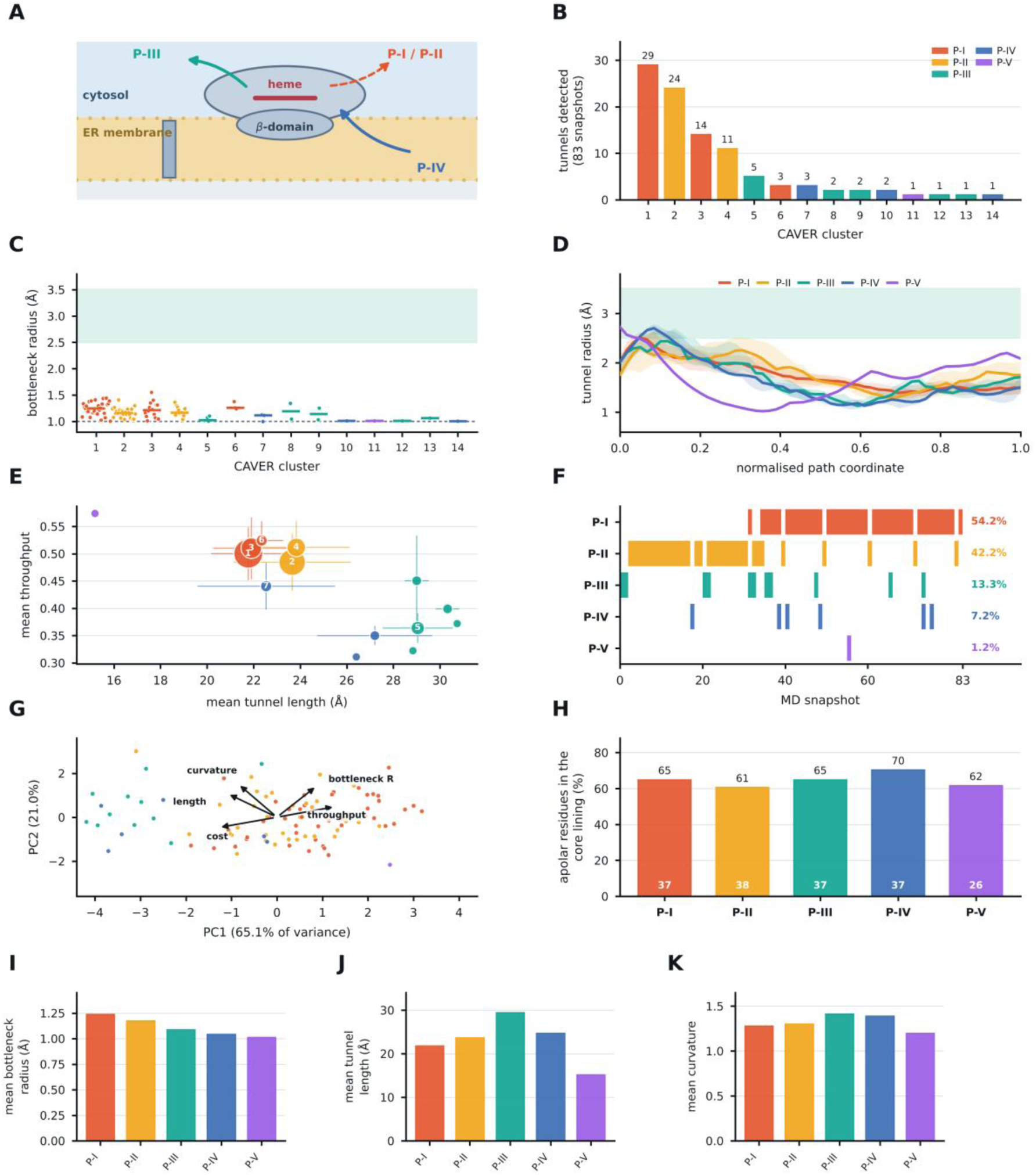
The CYP17A1 tunnel ensemble resolves into four constricted portals. (A) Catalytic domain anchored to the endoplasmic reticulum, with the four portals indicated. (B) Tunnels detected per CAVER cluster across 83 snapshots, colored by portal. (C) Bottleneck radius by cluster; points are individual tunnels and bars cluster medians. The shaded band marks the ∼2.5- 3.5 Å radius a steroid nucleus requires and the dotted line the 1.0 Å probe radius. (D) Median tunnel-radius profile of each portal along the normalized path coordinate, interquartile ranges shaded. (E) Mean length against mean throughput per cluster; bubble area is proportional to tunnel number. (F) Snapshots in which each portal was open. (G) Principal components of the five standardized tunnel descriptors, loadings shown as arrows. (H) Apolar fraction of the core lining of each portal, with the number of core lining residues inside each bar. (I-K) Mean bottleneck radius, length and curvature by portal. Source data: Supplementary Tables S1-S3 and Supplementary Figure S1.

Generally no tunnel is open permanently as observed. Bottleneck radii span 1.00-1.55 Å across all 99 tunnels, and every portal narrows from a wide active-site chamber to a mid-path constriction (Figure 1, Supplementary Table S3). Every portal therefore, must widen by side-chain rotation and backbone movements before a steroid can pass, consistent with the finding that P450 ligand tunnels are gated [25, 26, 35].

### 3.2. Residue architecture and the design of five ligand portal variants

Tunnel-lining analysis gives each portal a distinct signature (Figure 2, Supplementary Table S4). P-I and P-II share a core of B’-helix and B-C loop residues (Ile104, Ala105, Ser106, Asn107, Gly111, Phe114) with G-helix (Val236, Arg239, Asn240) and I-helix (Leu293, Thr294, Gly297, Asp298) contributions. P-III is lined by F-helix and F/G-loop residues (Tyr201, Asn202, Ile205, Ile206, Leu209) and by G-helix residues Leu242, Leu243 and Ile246. P-IV is lined by the β1-1 strand, the K’-β1-4 region (Pro365, Val366, Ala367, Met369, Ile371) and the β4 hairpin (Val482, Val483, Phe484). The tunnels share much of their core lining, reflecting a common active-site end rather than shared entry points, and all are strongly apolar there (61-70%), P-IV is the most hydrophobic, as expected for an interface facing the bilayer (Supplementary Table S6).

**Figure 2.**
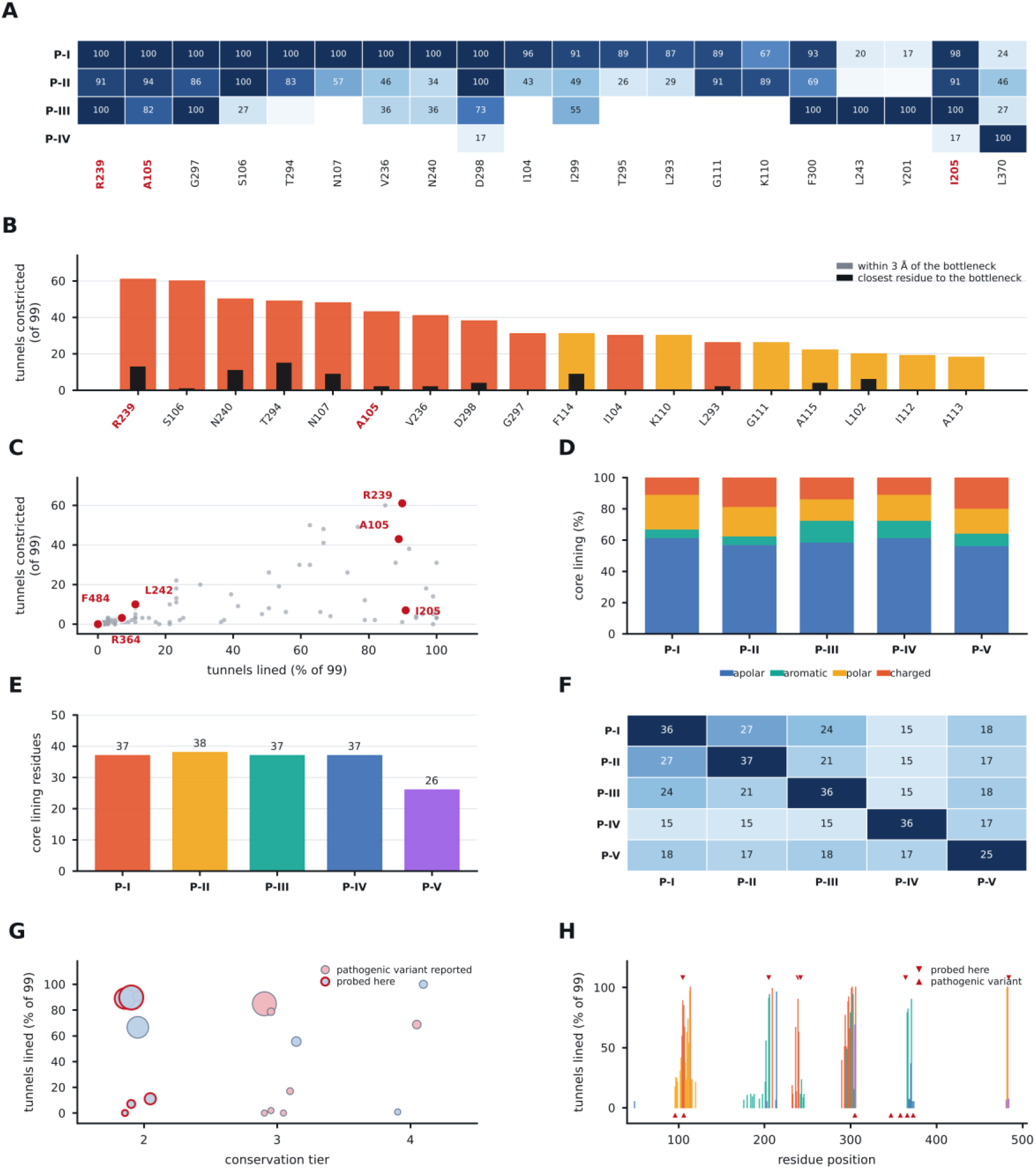
Residue-level architecture of the four portals. (A) Tunnel-lining occupancy of the residues that best discriminate between portals, as the percentage of each portal’s tunnels. (B) Bottleneck recurrence across all 99 tunnels; colored bars count observations within 3 Å of the bottleneck and black bars those in which the residue is closest to it. (C) Lining frequency against bottleneck recurrence for every residue. (D) Amino-acid class composition of each portal’s core lining, defined as residues present in at least 50% of its tunnels. (E) Size of each portal’s core lining. (F) Core lining residues shared between portal pairs. (G) Lining frequency against conservation tier; marker area scales with bottleneck recurrence. (H) Lining residues along the sequence. Positions probed by mutagenesis are marked in red throughout. Source data: Supplementary Tables S4-S8.

Bottleneck recurrence is far more selective than lining frequency, and the two are largely independent (Figure 2, Supplementary Table S5). Arg239 lies within 3 Å of the constriction in 61 of 99 tunnels, followed by Ser106 (60), Asn240 (50), Thr294 (49), Asn107 (48), Ala105 (43), Val236 (41) and Asp298 (38). Among the positions probed, Leu242 constricts 10 tunnels and Ile205 seven, both within P-III, while Phe484 constricts three, all within P-IV, where it is the bottleneck residue of every cluster-7 tunnel. Arg364 was not detected in this apo ensemble.

Five variants were constructed to study the details of substrate access (Supplementary Tables S7 and S8). A105L mutation reproduces the published probe of the dominant P-I/P-II bottleneck [15]. L242W mutation places the largest natural side chain in the P-III lumen, whereas I205C/R239C mutation introduces a cysteine pair across the P-III entry, the two positions lying on opposite sides of this tunnel. the R364W and R364C/F484C mutations target P-IV, by steric bulk at the loop preceding the β1-4 strand that forms the inner wall of the tunnel and by restraining the β1-4/β4 hairpin around Phe484.

### 3.3. The CYP17A1 Catalytic phenotypes separate the tunnels into two classes

Both CYP17A1 reactions were assayed in multiple independent experiments (Figure 3, Supplementary Table S9, Supplementary Figure S2), and the panel separates into two classes whose mean activities differ by nearly two orders of magnitude, with no mutation in between. A105L mutation, which places a leucine at the most frequently sampled bottleneck of the ensemble, exceeded wild type activity: 116.1 ± 8.0% of wild-type 17α-hydroxylase activity (95% CI 107.8-124.5, P = 0.0043) and 134.8 ± 27.9% of 17,20-lyase activity (95% CI 105.5-164.1, P = 0.028; Supplementary Table S10), reproducing the mild increase previously reported for the same substitution [15]. The Ala105 is in 43 of 99 tunnels and P-I plus P-II account for 82% of the ensemble, yet adding a methylene-rich side chain there does not impact the enzyme, indicating sampling frequency is a poor proxy for steroid flux [25].

**Figure 3.**
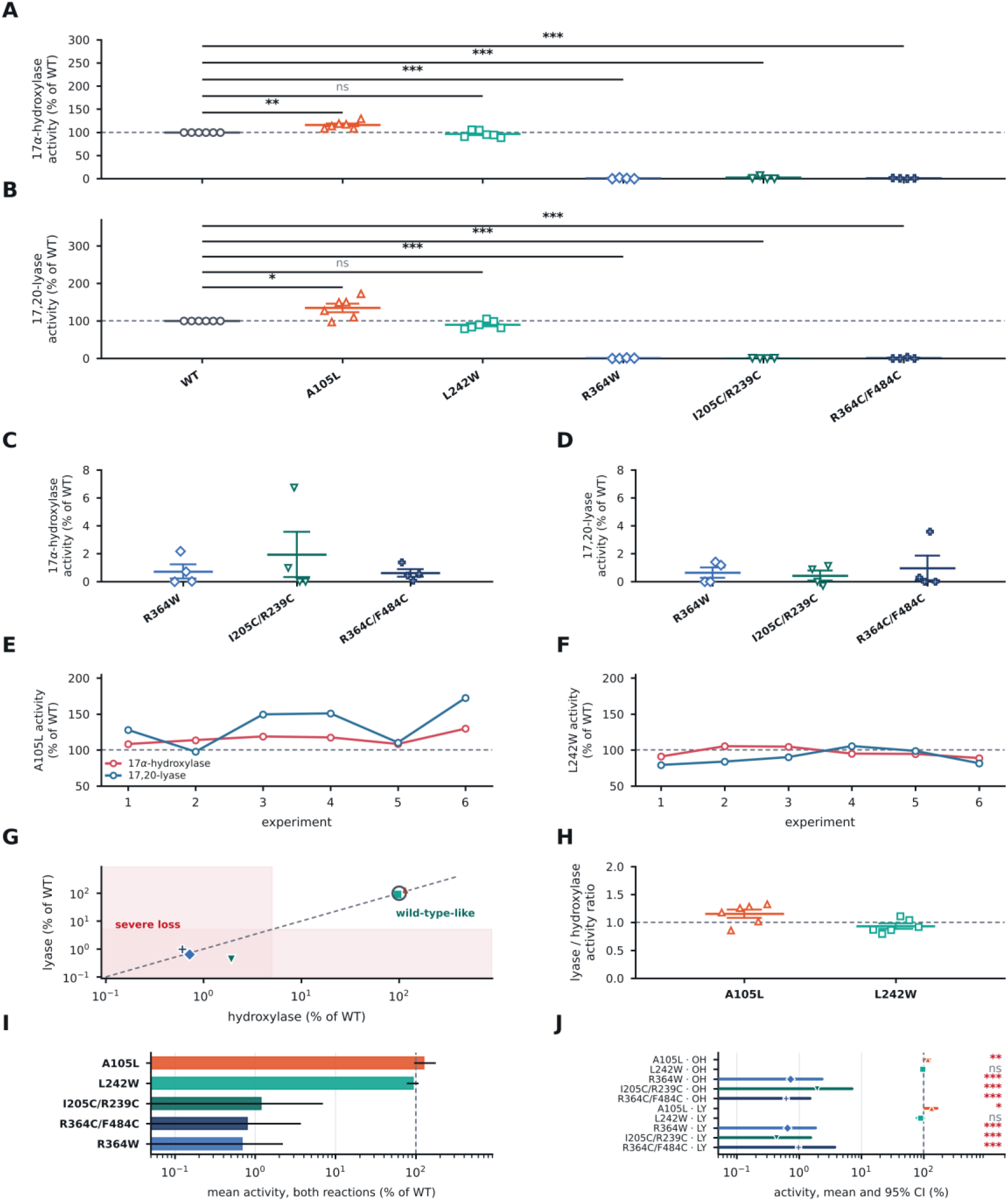
Catalytic phenotypes of the portal variants. (A, B) 17α-Hydroxylase (A) and 17,20-lyase (B) activity as a percentage of the wild-type control assayed in parallel. (C, D) The severe-loss class from A and B on a linear axis. (E, F) Activity of A105L (E) and L242W (F) in each experiment. (G) Phenotype space built from the variant means; the diagonal marks equal effect on both reactions. (H) Lyase-to-hydroxylase activity ratio, undefined below 5% residual hydroxylase activity. (I) Severity ranking across both reactions on a log scale. (J) Effect estimate with its 95% confidence interval for every variant and reaction. Points are independent experiments, the bar the mean and whiskers the SEM; significance is from a two-sided one-sample t-test against wild type: ns, P ≥ 0.05; *, P < 0.05; **, P < 0.01; ***, P < 0.001. Source data: Supplementary Tables S9 and S10 and Supplementary Figure S2.

Mutation L242W, a tryptophan in the middle of the P-III lumen, was equally well tolerated: 96.6 ± 6.9% hydroxylase activity (P = 0.28) and 89.9 ± 10.4% lyase activity (P = 0.063). Given a wild-type bottleneck of only 1.09 Å and an indole adding roughly 40 Å³ of volume, a rigid-pore model predicts near-complete blockade. The tolerance indicates that the lumen accommodates the substitution by local rearrangements for a steroid to pass.

The I205C/R239C mutation, which constrains the same tunnel across its entry, abolished CYP17A1 activities: 1.9 ± 3.2% hydroxylase and 0.4 ± 0.7% lyase activity (both P < 0.001). Therefore, adding bulk inside the tunnel is tolerated but restraining the relative motion of the F and G helices is not. Arg239 lies at the most recurrent constriction of the ensemble and Ile205 lines 91% of all tunnels, so the pair indicates the mobility of the region forming the upper part of the active site. P-III therefore behaves as a conformationally gated exit rather than a static pore. Cytochrome b₅ may stimulate the lyase reaction without necessarily transferring electrons [19, 20, 36], and NMR places its contacts and the associated conformational changes in the same F/G region [37, 38], so part of cytochrome b5 mediated stimulation may be mechanical.

Both perturbations of the membrane-facing P-IV portal produced near-complete loss of function. R364W reduced activity to 0.7 ± 1.0% (hydroxylase) and 0.6 ± 0.8% (lyase) of wild type, and R364C/F484C to 0.6 ± 0.6% and 1.0 ± 1.7% (all P < 0.001). The two agree closely despite acting by different mechanisms, and both reactions are affected equally, as expected upstream of the branch point: if substrate cannot reach the heme, neither reaction can proceed. P-IV is the only portal whose entry lies on the membrane-facing β-domain surface, which simulation and nanodisc work of Sligar and colleagues place in contact with the bilayer [17, 18], and it is the most hydrophobic of the four in its core lining, consistent with P-IV as the obligate ingress portal drawing substrate directly from the lipid phase.

Val366 sits inside this tunnel past its constriction and lines 79% of all tunnels. The clinical V366M variant causes isolated 17,20-lyase deficiency, simulations showing the methionine projecting its methyl group toward the heme iron and increasing the mean O17-Fe distance for bound 17OH-pregnenolone [6]. Because the lyase reaction needs closer approach of 17α-hydroxylated substrate to reactive iron-oxygen species [39], a modest steric intrusion disables it while leaving hydroxylation reaction intact. The gating portals control which molecules reach the site and the Val366 environment controls how closely substrates reach near the iron.

### 3.4. CYP17A1 protein levels do not account for the phenotypes

CYP17A1 abundance was at wild-type level for the variants that retain activity (A105L 124% and 126% of wild type; L242W 124% and 124%) and 31-57% of wild type for the three inactive variants (Figure 4, Supplementary Table S9). Measured activity is reported as the primary readout, with protein abundance alongside as a qualitative control and the expression-corrected values are given in Figure 4 and Supplementary Tables S10 and S11. The tolerated variants accumulate at or above wild type, so their measured activity has a lower impact on their specific activity, while the inactive variants accumulate at a third to a half of wild type, so even the most generous correction leaves them impaired more than twenty-fold (Supplementary Table S12).

**Figure 4.**
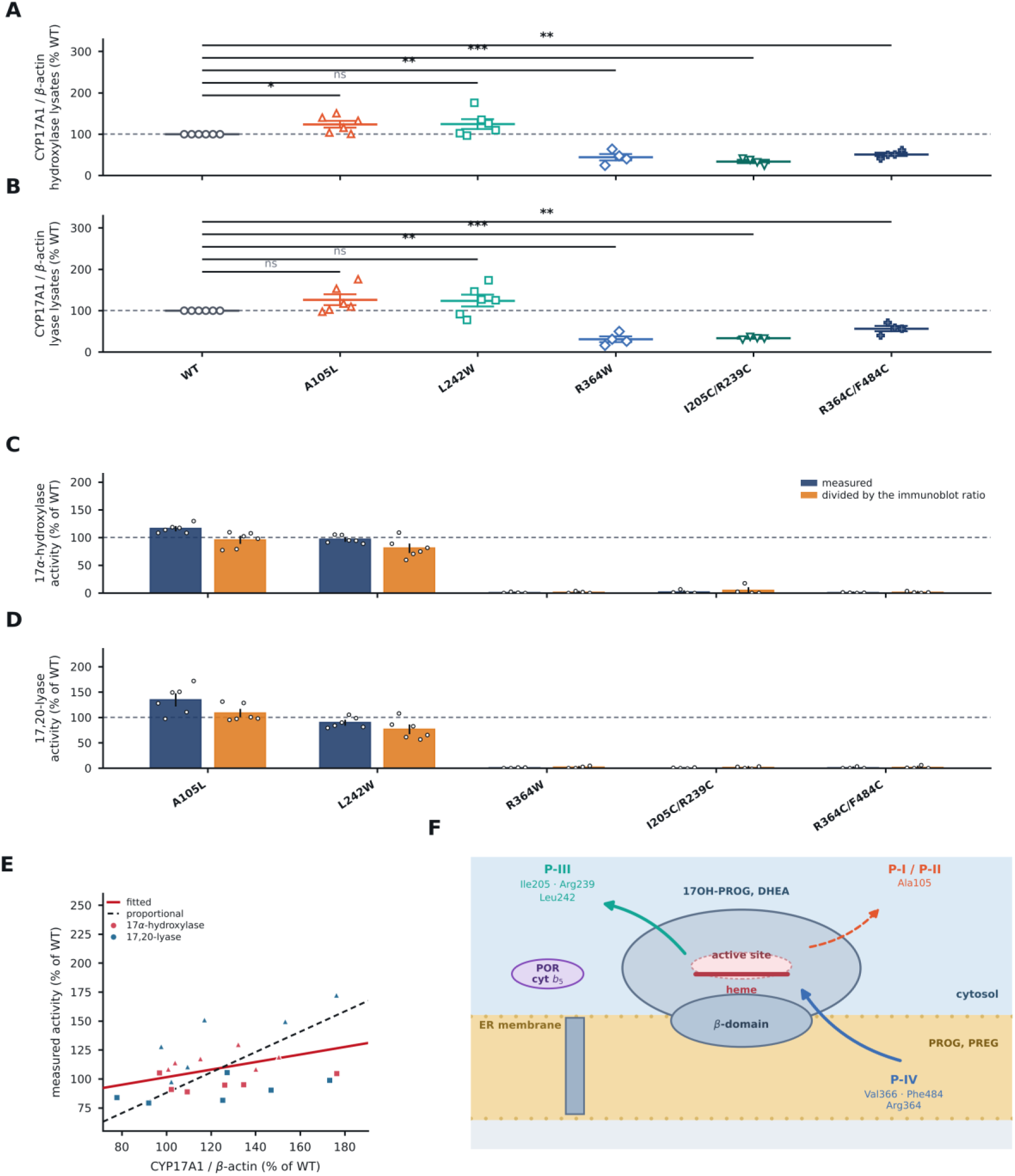
CYP17A1 protein levels and the dual-portal model. (A, B) Immunoblot CYP17A1/β-actin ratio of the hydroxylase-assay (A) and lyase-assay (B) lysates, as a percentage of wild type. (C, D) Measured and expression-corrected 17α-hydroxylase (C) and 17,20-lyase (D) activity. (E) Measured activity against the matched immunoblot ratio for the two variants that retain activity. (F) Dual-portal model: substrate enters from the bilayer through P-IV past Val366 and Phe484, product leaves through the conformationally gated F/G portal P-III, and P-I and P-II, dominant in the apo ensemble, are not rate-limiting. Points are independent experiments, the bar the mean and whiskers the SEM. Source data: Supplementary Tables S9-S12 and Supplementary Figure S3.

### 3.5. A dual-portal model of vectorial ligand transport in CYP17A1

Combining tunnel geometry, portal-specific phenotypes and membrane topology gives the model presented in Figure 4: substrate is taken up from the bilayer through P-IV past Phe484 and the Val366 gate, catalysis proceeds in an enclosed site whose steric properties around Val366 discriminate between the two substrates, and product is released to the cytosol through P-III, which opens by F/G breathing rather than by presenting a fixed lumen. P-I and P-II, though dominant in the apo trajectory, are not rate-limiting.

These data support directional selectivity in steroid access. The portals face opposite sides of the CYP17A1, and the phenotypes differ in kind with P-IV perturbations abolishing both reactions equally while P-III perturbations tolerated adding bulk, but removing motion was not tolerated. Based on our studies two inhibitor design strategies can be suggested. A compound binding the P-IV entry would inhibit all CYP17A1 chemistry non-competitively, matching R364W mutation. More importantly, a compound restricting F/G motion without occluding the active site should impair the reaction with the stricter geometric constraint on lyase activity, while minimally impacting hydroxylase activity, which is ideal in a prostate-cancer drug needs [10]. A secondary binding site in CYP17A1 found by enhanced-sampling simulation lies in a comparable peripheral region [40].

The tunnel analysis enumerates candidate tunnels but cannot estimate their flux [25]. Arg364 did not appear as a lining or bottleneck residue of any tunnel, so the mechanism of R364W remains open; Phe484, by contrast, constricts every cluster-7 tunnel, so the P-IV assignment does not depend on Arg364. Human CYP17A1 presents at least four geometrically distinct tunnels between its buried active site and the surface, none allowing access to steroids in resting states. Functional probing indicated that two most frequently sampled tunnels are dispensable, the F/G tunnel is required in its capacity to move rather than as a fixed lumen, and the membrane-facing β-domain tunnel is definitely required.

## 4. Conclusions

Human CYP17A1 presents at least four geometrically distinct paths between its buried active site and the protein surface, and none of them is open for a steroid in the resting state. Our functional analysis separated these routes. The two most frequently sampled routes of the apo ensemble, P-I and P-II, are dispensable: a leucine at their shared bottleneck leaves both CYP17A1 reactions intact, so sampling frequency is a poor proxy for catalytic relevance. The F/G route, P-III, accommodates a tryptophan in its lumen but not a cysteine pair restraining helix motion, so this portal is required for its capacity to move rather than as a fixed channel. The membrane-facing β-domain route, P-IV, is required: two chemically distinct perturbations, added bulk at Arg364 and a constrained β1-4/β4 hairpin at Phe484, each reduced both CYP17A1 reactions to below 2% of wild type, well beyond what the accompanying reduction in enzyme accumulation can explain.

Together these observations support a dual-portal model of vectorial ligand transport, in which CYP17A1 draws lipophilic substrate directly from the bilayer through P-IV and releases the more polar product to the cytosol through a conformationally gated P-III. The model assigns a mechanical role to the F/G region that carries the cytochrome b₅ contacts and the ligand-dependent conformational changes reported by NMR, and places steric discrimination between the two substrates at Val366, inside the ingress route. Our model also identifies two peripheral surfaces for inhibitor design: the P-IV entry, where a compound would block all CYP17A1 chemistry, and the F/G region, where restricting the dynamic motion may impair the CYP17A1 lyase activity selectively.

## Data availability

Supplementary Data (Supplementary_Data.xlsx) contains Supplementary Tables S1-S12, itemized in its contents sheet: tunnel and portal geometry (S1-S3), residue architecture, annotation and variant design (S4-S8), activity and immunoblot data with the variant summary statistics (S9, S10) and the expression-corrected analysis (S11, S12). Supplementary Materials (Supplementary_Materials.pdf) contains the Supplementary Methods and Supplementary Figures S1-S3. The primary CAVER output is provided as tunnel_characteristics.csv, tunnel_profiles.csv, bottlenecks.csv, residues.txt and ids_and_clusters.csv.

## Acknowledgements

Supported by Cancer Research Switzerland (KFS 5557-02-2022),

## Author contributions

**CRediT:** Conceptualization, A.V.P.; Data curation, Y.X.; A.Z.; Funding acquisition, A.V.P.; Investigation, Y.X., A.Z.; Methodology, Y.X., A.Z., A.V.P.; Project administration, A.V.P.; Supervision, A.V.P.; Validation, Y.X., A.Z., A.V.P.; Visualization, Y.X., A.Z., A.V.P.; Writing – original draft, Y.X., A.V.P.; Writing – review and editing, Y.X., A.Z., A.V.P.

## References

1. Miller, W.L. and R.J. Auchus, The molecular biology, biochemistry, and physiology of human steroidogenesis and its disorders. Endocr Rev, 2011. 32(1): p. 81–151.

2. Mak, P.J., et al., Human Cytochrome CYP17A1: The Structural Basis for Compromised Lyase Activity with 17-Hydroxyprogesterone. J Am Chem Soc, 2018. 140(23): p. 7324– 7331.

3. Yanase, T., et al., *Combined 17 alpha-hydroxylase/17,20-lyase deficiency due to a 7-basepair duplication in the N-terminal region of the cytochrome P45017 alpha (CYP17)* gene. J Clin Endocrinol Metab, 1990. 70(5): p. 1325–9.

4. Geller, D.H., et al., The genetic and functional basis of isolated 17,20-lyase deficiency. Nat Genet, 1997. 17(2): p. 201–5.

5. Sherbet, D.P., et al., CYP17 mutation E305G causes isolated 17,20-lyase deficiency by selectively altering substrate binding. J Biol Chem, 2003. 278(49): p. 48563–9.

6. Fernandez-Cancio, M., et al., Mechanism of the Dual Activities of Human CYP17A1 and Binding to Anti-Prostate Cancer Drug Abiraterone Revealed by a Novel V366M Mutation Causing 17,20 Lyase Deficiency. Pharmaceuticals (Basel), 2018. 11(2): p. 37.

7. de Bono, J.S., et al., Abiraterone and increased survival in metastatic prostate cancer. N Engl J Med, 2011. 364(21): p. 1995–2005.

8. Attard, G., et al., *Phase I clinical trial of a selective inhibitor of CYP17, abiraterone acetate, confirms that castration-resistant prostate cancer commonly remains hormone* driven. J Clin Oncol, 2008. 26(28): p. 4563–71.

9. Yakubu, J., T. du Toit, and A.V. Pandey, Multi-Omics Profiling Exposes a Metabolic Void in Prostate Cancer Models That Drives Precursor Dependence and Clinical Progression, in Preprints. 2026, Preprints.

10. Wrobel, T.M., et al., Non-steroidal CYP17A1 Inhibitors: Discovery and Assessment. J Med Chem, 2023. 66(10): p. 6542–6566.

11. Udhane, S.S., et al., Specificity of anti-prostate cancer CYP17A1 inhibitors on androgen biosynthesis. Biochem Biophys Res Commun, 2016. 477(4): p. 1005–1010.

12. Malikova, J., et al., CYP17A1 inhibitor abiraterone, an anti-prostate cancer drug, also inhibits the 21-hydroxylase activity of CYP21A2. J Steroid Biochem Mol Biol, 2017. 174: p. 192–200.

13. DeVore, N.M. and E.E. Scott, Structures of cytochrome P450 17A1 with prostate cancer drugs abiraterone and TOK-001. Nature, 2012. 482(7383): p. 116–9.

14. Petrunak, E.M., et al., Structural and Functional Evaluation of Clinically Relevant Inhibitors of Steroidogenic Cytochrome P450 17A1. Drug Metab Dispos, 2017. 45(6): p. 635–645.

15. Petrunak, E.M., et al., Structures of human steroidogenic cytochrome P450 17A1 with substrates. J Biol Chem, 2014. 289(47): p. 32952–64.

16. Burris-Hiday, S.D. and E.E. Scott, Steroidogenic cytochrome P450 17A1 structure and function. Mol Cell Endocrinol, 2021. 528: p. 111261.

17. Baylon, J.L., et al., Characterizing the membrane-bound state of cytochrome P450 3A4: structure, depth of insertion, and orientation. J Am Chem Soc, 2013. 135(23): p. 8542–51.

18. Denisov, I.G. and S.G. Sligar, Nanodiscs in Membrane Biochemistry and Biophysics. Chem Rev, 2017. 117(6): p. 4669–4713.

19. Auchus, R.J., T.C. Lee, and W.L. Miller, Cytochrome b5 augments the 17,20-lyase activity of human P450c17 without direct electron transfer. J Biol Chem, 1998. 273(6): p. 3158–65.

20. Pandey, A.V. and W.L. Miller, Regulation of 17,20 lyase activity by cytochrome b5 and by serine phosphorylation of P450c17. J Biol Chem, 2005. 280(14): p. 13265–71.

21. Miller, W.L., A.V. Pandey, and C.E. Fluck, Disordered Electron Transfer: New Forms of Defective Steroidogenesis and Mitochondriopathy. J Clin Endocrinol Metab, 2025. 110(3): p. e574–e582.

22. Pandey, A.V. and C.E. Flück, NADPH P450 oxidoreductase: structure, function, and pathology of diseases. Pharmacology & therapeutics, 2013.

23. Khatri, Y., et al., Active site proton delivery and the lyase activity of human CYP17A1. Biochem Biophys Res Commun, 2014. 443(1): p. 179–84.

24. Liu, Y., et al., Substrate-Specific Allosteric Effects on the Enhancement of CYP17A1 Lyase Efficiency by Cytochrome b(5). J Am Chem Soc, 2021. 143(10): p. 3729–3733.

25. Cojocaru, V., P.J. Winn, and R.C. Wade, The ins and outs of cytochrome P450s. Biochim Biophys Acta, 2007. 1770(3): p. 390–401.

26. Urban, P., et al., Ligand Access Channels in Cytochrome P450 Enzymes: A Review. Int J Mol Sci, 2018. 19(6): p. 1617.

27. Krieger, E. and G. Vriend, New ways to boost molecular dynamics simulations. J Comput Chem, 2015. 36(13): p. 996–1007.

28. Fernandez-Cancio, M., et al., Mechanism of the Dual Activities of Human CYP17A1 and Binding to Anti-Prostate Cancer Drug Abiraterone Revealed by a Novel V366M Mutation Causing 17,20 Lyase Deficiency. Pharmaceuticals (Basel), 2018. 11(2).

29. Chovancova, E., et al., CAVER 3.0: a tool for the analysis of transport pathways in dynamic protein structures. PLoS Comput Biol, 2012. 8(10): p. e1002708.

30. Dombkowski, A.A., K.Z. Sultana, and D.B. Craig, Protein disulfide engineering. FEBS Lett, 2014. 588(2): p. 206–12.

31. Xu, Y., et al., Integrated mapping resolves pathogenicity of 11 CYP21A2 variants in congenital adrenal hyperplasia. J Endocr Soc, 2026. 10(7): p. bvag127.

32. Yakubu, J., et al., Nanoparticles with curcumin and piperine modulate steroid biosynthesis in prostate cancer. Sci Rep, 2025. 15(1): p. 13613.

33. Grudzińska, A., et al., Discovery and Characterization of Novel Piperidine Derivatives as Potent Cytochrome P450 17A1 Inhibitors for Prostate Cancer Therapy. ChemMedChem, 2026. 21(19): p. e70336.

34. Wrobel, T.M., et al., Pyridine indole hybrids as novel potent CYP17A1 inhibitors. J Enzyme Inhib Med Chem, 2025. 40(1): p. 2463014.

35. Bouchoucha, N., et al., Characterization of a novel CYP19A1 (aromatase) R192H mutation causing virilization of a 46,XX newborn, undervirilization of the 46,XY brother, but no virilization of the mother during pregnancies. Mol Cell Endocrinol, 2014. 390(1- 2): p. 8–17.

36. Duggal, R., et al., Evidence that cytochrome b5 acts as a redox donor in CYP17A1 mediated androgen synthesis. Biochem Biophys Res Commun, 2016. 477(2): p. 202–8.

37. Estrada, D.F., J.S. Laurence, and E.E. Scott, Substrate-modulated cytochrome P450 17A1 and cytochrome b5 interactions revealed by NMR. J Biol Chem, 2013. 288(23): p. 17008–17018.

38. Estrada, D.F., et al., Human cytochrome P450 17A1 conformational selection: modulation by ligand and cytochrome b5. J Biol Chem, 2014. 289(20): p. 14310–20.

39. Gonzalez, E. and F.P. Guengerich, Kinetic processivity of the two-step oxidations of progesterone and pregnenolone to androgens by human cytochrome P450 17A1. J Biol Chem, 2017. 292(32): p. 13168–13185.

40. Wrobel, T.M., D. Bartuzi, and A.A. Kaczor, Secondary Binding Site of CYP17A1 in Enhanced Sampling Simulations. J Chem Inf Model, 2024. 64(19): p. 7679–7686.

