## Supplementary Materials for "Dual-portal ligand transport in human cytochrome P450 17A1"

### Contents

Supplementary Methods 1. Structure preparation
Supplementary Methods 2. Molecular dynamics
Supplementary Methods 3. Tunnel detection, clustering and portal assignment
Supplementary Methods 4. Tunnel descriptors and principal-component analysis
Supplementary Methods 5. Lining residues, bottleneck recurrence and lining composition
Supplementary Methods 6. Disulfide design and variant selection
Supplementary Methods 7. Mutagenesis, expression and activity assays
Supplementary Methods 8. Immunoblotting
Supplementary Methods 9. Statistics
Supplementary Methods 10. Conservation and clinical-variant annotation
Supplementary Methods 11. Figure preparation, source data and code
Supplementary Figures S1-S3
Supplementary Tables S1-S12 (provided as Supplementary Data, .xlsx)
Supplementary references

### Supplementary Methods 1. Structure preparation

The catalytic domain of human CYP17A1 was taken from the abiraterone complex (PDB 3RUK, chain D), selected for the smallest number of unmodelled residues, with cross-reference to the substrate complexes 4NKX and 4NKZ and to 3SWZ [1-3]. Residues preceding Arg45 were removed: this segment is largely unresolved in the available structures and lacks ordered secondary structure. A single ordered water molecule adjacent to the heme and position 366 was retained because it is conserved across the available CYP17A1 structures and lies at the inner end of the β-domain route. Hydrogens were added at pH 7.4 and the structure was energy-minimised before simulation. The orientation of the catalytic domain relative to the bilayer follows the consensus for microsomal P450s [4].

**Supplementary Methods 2. Molecular dynamics**

Simulations of the wild-type apo enzyme were run at 310 K in explicit solvent as previously described [5]. Snapshots were written at regular intervals along the trajectory and converted to PDB format; 83 snapshots entered the tunnel analysis reported here. Trajectory processing, superposition and visual inspection were performed in UCSF Chimera [6].

**Supplementary Methods 3. Tunnel detection, clustering and portal assignment**

Tunnels were computed with the command-line version of CAVER 3.0 using a probe radius of 1.0 Å, with the starting point defined at the heme iron. Clustering across snapshots used the default CAVER weighting. The calculation returned 99 transport pathways distributed over 14 clusters across the 83 snapshots.

The 14 clusters are not independent: several share an origin at the active site and diverge only near the protein surface. Portals were therefore assigned by the secondary-structure elements that form the tunnel mouth, using the lining-residue lists; clusters whose surface openings were bounded by the same elements were grouped. This gives P-I (clusters 1, 3, 6), P-II (clusters 2, 4), P-III (clusters 5, 8, 9, 12, 13), P-IV (clusters 7, 10, 14) and the singleton P-V (cluster 11). Assignment to the established pathway nomenclature [7, 8] was made by comparing the lining-residue sets with the published pathway definitions. Per-cluster and per-portal statistics are in Supplementary Tables S1 and S2.

**Supplementary Methods 4. Tunnel descriptors and principal-component analysis**

Five per-tunnel descriptors were exported from CAVER: throughput, cost, bottleneck radius, length and curvature. Radius profiles were interpolated onto a common normalised path coordinate of 60 points running from the active site (0) to bulk solvent [9] before computing medians and interquartile ranges, so that tunnels of different length could be compared (Supplementary Table S3).

For the principal-component analysis the five descriptors were centred and scaled to unit variance across all 99 tunnels, the covariance matrix was diagonalised by Jacobi rotation, and each tunnel was projected onto the first two components. PC1 accounts for 65.1% and PC2 for 21.0% of the variance. PC1 contrasts short, wide, high-throughput tunnels (positive loadings on throughput and bottleneck radius) with long, curved, narrow ones (positive loadings on cost, length and curvature).

**Supplementary Methods 5. Lining residues, bottleneck recurrence and lining composition**

Tunnel-lining residues were taken as those within 3.0 Å of the tunnel axis, as reported by CAVER in residues.txt. Occupancy is expressed as the percentage of a portal's tunnels lined by a given residue, capped at the number of tunnels in the contributing cluster (Supplementary Table S4).

Bottleneck recurrence was taken from bottlenecks.csv, which lists the residues within 3.0 Å of the bottleneck of each tunnel, ordered from the closest. Two counts are reported: the number of tunnels in which a residue lies within 3 Å of the bottleneck, and the number in which it is the residue closest to it (Supplementary Table S5).

The core lining of a portal was defined as the residues present in at least 50% of that portal's tunnels. Composition was computed over this set, classifying residues as apolar (AVLIMPGC), aromatic (FWY), polar (STNQHC) or charged (DEKR); the apolar fraction quoted in the main text treats aromatics as apolar, consistent with the convention used for ligand tunnels (Supplementary Table S6).

**Supplementary Methods 6. Disulfide design and variant selection**

Candidate cysteine pairs were evaluated by Cβ-Cβ distance and side-chain geometry in the crystal structures and in MD snapshots, and independently with the Disulfide by Design approach. Homology models carrying each pair were built using 3SWZ, 3RUK and 4NKZ as forced templates and inspected for secondary-structure distortion and for bond formation. Neither the I205C/R239C nor the R364C/F484C pair was predicted to form a disulfide spontaneously despite the residues being in proximity, so these variants may act by introducing unpaired cysteines and perturbing local packing rather than by cross-linking.

The five variants and the reasoning behind each are listed in Supplementary Table S8.

**Supplementary Methods 7. Mutagenesis, expression and activity assays**

Variants were generated in a human CYP17A1 expression construct by site-directed mutagenesis and verified by Sanger sequencing across the full coding region. HEK293 cells were transfected transiently and harvested 48 h after transfection.

17α-Hydroxylase activity was measured as conversion of progesterone to 17α-hydroxyprogesterone, and 17,20-lyase activity as conversion of 17α-hydroxypregnenolone to dehydroepiandrosterone. Products were resolved by thin-layer chromatography and quantified by phosphorimaging. Each experiment included technical replicates and wild-type, empty-vector and untransfected controls, and activities are reported as a percentage of the wild-type control assayed in parallel on the same plate in the same experiment. Variants retaining measurable activity were carried through further rounds of assay, so more data points are available for them. Per-experiment values, including the technical-replicate standard deviations, are in Supplementary Table S9 and Supplementary Figure S2.

Authors: before submission, insert the expression vector, mutagenesis kit, transfection reagent and cell numbers, the substrate concentrations and specific activities, the incubation time and temperature, the extraction procedure, the TLC solvent system, and the imaging and quantification software.

**Supplementary Methods 8. Immunoblotting**

CYP17A1 abundance was determined by immunoblot as previously described in the same lysates used for the activity assays, with β-actin as loading control, and is expressed as the CYP17A1/β-actin ratio relative to the wild-type lane of the same membrane [10]. Separate lysate sets were blotted for the hydroxylase and lyase assays, so each variant has two abundance values per experiment (Supplementary Table S9).

Expression-corrected activity was calculated as measured activity (% of wild type) divided by CYP17A1 abundance (% of wild type), × 100, using the abundance of the matching lysate (Supplementary Tables S11 and S12).

**Supplementary Methods 9. Statistics**

Variant-level data are presented as the mean ± SD of independent experiments, with the individual experiments overlaid on each panel. Each variant was compared with the wild-type reference, defined as 100% in every experiment, by a two-sided one-sample t-test; significance is indicated as ns ($P \geq 0.05$), * ($P < 0.05$), ** ($P < 0.01$) and *** ($P < 0.001$). Confidence intervals are 95% intervals of the mean from the same t distribution. Tests were run on measured activity and on expression-corrected activity; the classification into wild-type-like and severe-loss classes is identical in both (Supplementary Tables S10 and S12). All statistics were computed in Python 3.12 using the implementations in activity_analysis.py, which are self-contained and do not depend on external statistical packages.

**Supplementary Methods 10. Conservation and clinical-variant annotation**

Positions were annotated for secondary-structure element, substrate-recognition-site membership [11], tunnel-lining frequency and bottleneck recurrence (from the CAVER output above), and for reported pathogenic variants from the published 17α-hydroxylase deficiency and isolated 17,20-lyase deficiency literature (Supplementary Table S7).

Conservation tiers are qualitative assignments on a four-point scale: tier 4, invariant across the P450 superfamily (the heme-ligating cysteine and the I-helix acid-alcohol pair); tier 3, strongly conserved in vertebrate CYP17A1 and related steroidogenic P450s; tier 2, conserved among CYP17A1 orthologues; tier 1, variable. They are knowledge-based assignments derived from P450 motif architecture and the published mutational literature, adequate for the qualitative contrast drawn in the main text: catalytic positions are invariant, transport positions are orthologue-level.

**Supplementary Methods 11. Figure preparation, source data and code**

Figures were generated in Python 3.12 with Matplotlib directly from the primary CAVER output and the primary activity data.

The primary CAVER output is provided as tunnel_characteristics.csv, tunnel_profiles.csv, bottlenecks.csv, residues.txt and ids_and_clusters.csv.

Per-panel source data are provided as ten CSV files: SD_Fig1_cluster_statistics.csv, SD_Fig1_portal_summary.csv, SD_Fig1_radius_profiles.csv, SD_Fig2_bottleneck_recurrence.csv, SD_Fig2_lining_occupancy.csv, SD_Fig2_portal_hydrophobicity.csv, SD_Fig3_per_experiment.csv, SD_Fig3_variant_summary.csv, SD_Fig4_normalisation.csv and SD_Fig4_precision.csv.

The analysis and figure code consists of activity_data.py (the primary activity and immunoblot values, transcribed from the laboratory workbook), activity_analysis.py, tunnel_analysis.py, residue_annotation.py, figstyle.py, fig1_tunnels.py, fig2_residues.py, fig3_activity.py, fig4_model.py, graphical_abstract.py, supp_figures.py, build_supplementary_data.py and build_documents.py.

### Supplementary Figures

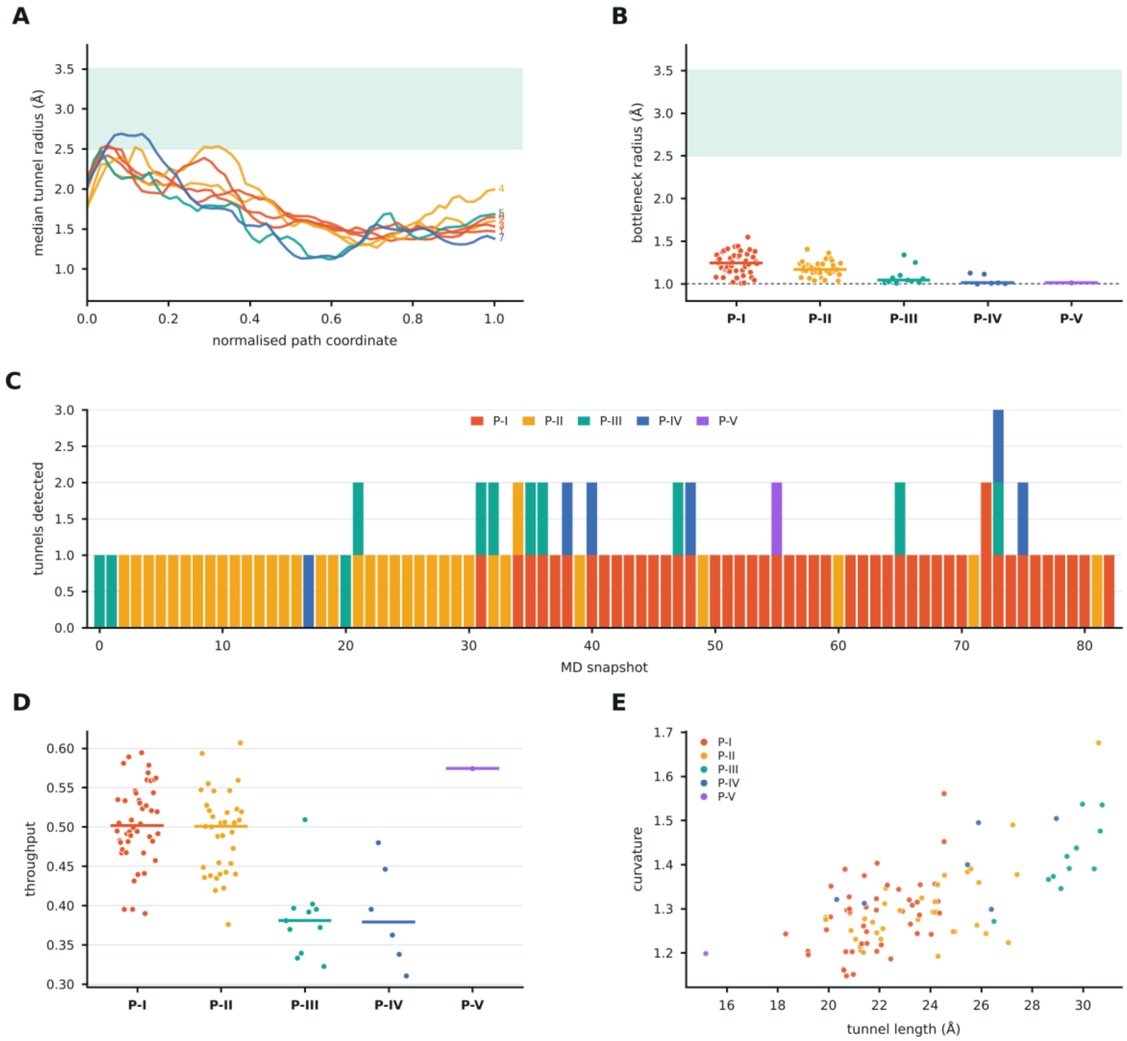


**Supplementary Figure S1.** Geometry of the CYP17A1 apo tunnel ensemble. (A) Median radius profile of each well-sampled CAVER cluster along the normalized path coordinate, colored by portal and labelled with the cluster number. (B) Bottleneck radii of all 99 tunnels grouped by portal; points are individual tunnels and bars portal medians. The shaded band marks the ~2.5-3.5 Å radius required by a steroid nucleus and the dotted line the 1.0 Å CAVER probe radius. (C) Tunnels detected in each of the 83 snapshots, stacked by portal. (D) Throughput of all tunnels by portal. (E) Tunnel length against curvature for all 99 tunnels, colored by portal. Source data: Supplementary Tables S1-S3.

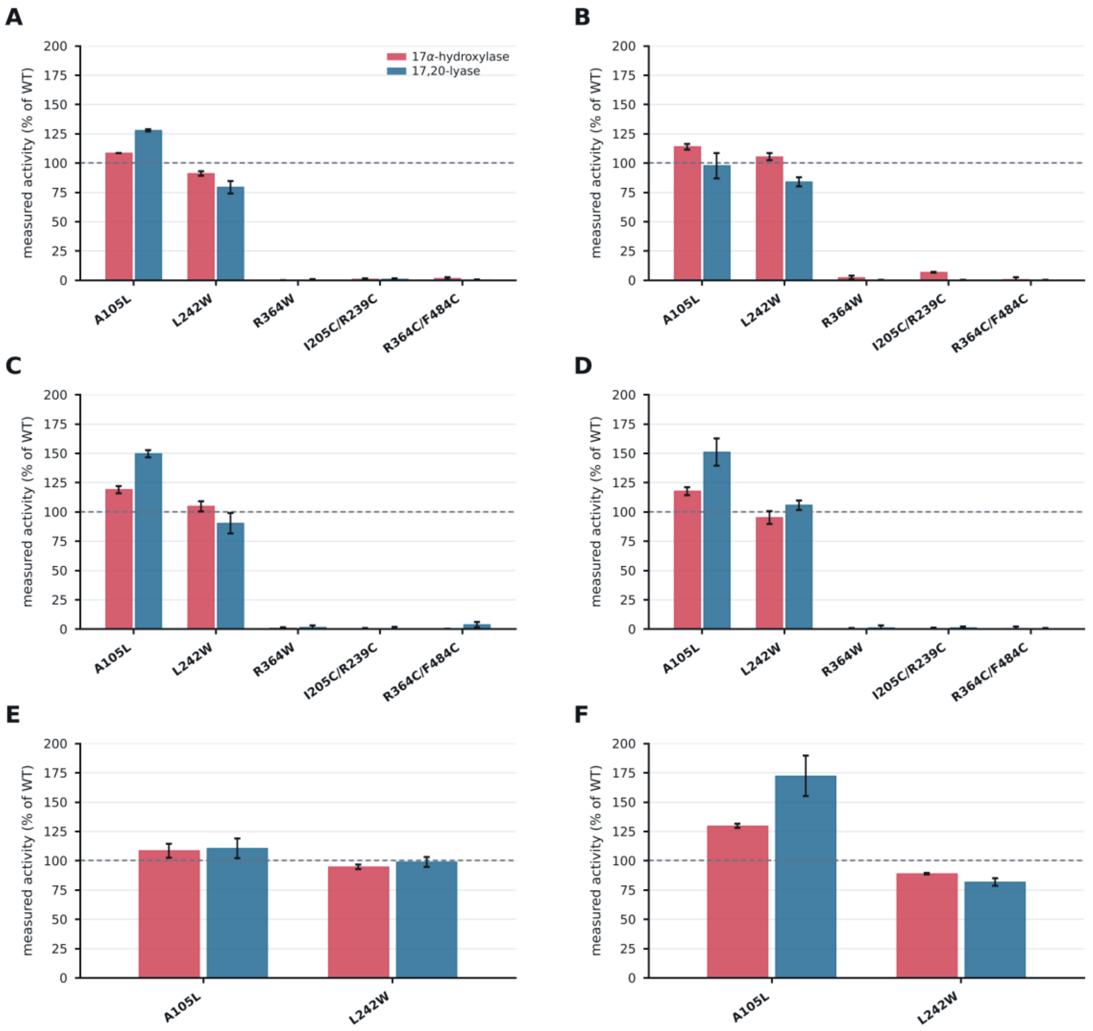


**Supplementary Figure S2.** Measured activity in each independent experiment. (A-F) One experiment per panel, showing 17α-hydroxylase and 17,20-lyase activity for every construct assayed in that experiment, expressed as a percentage of the wild-type control assayed in parallel. Bars show the mean of the technical replicates within the experiment and whiskers their standard deviation. The partition into a wild-type-like class and a severe-loss class is reproduced in every experiment. Source data: Supplementary Table S9.

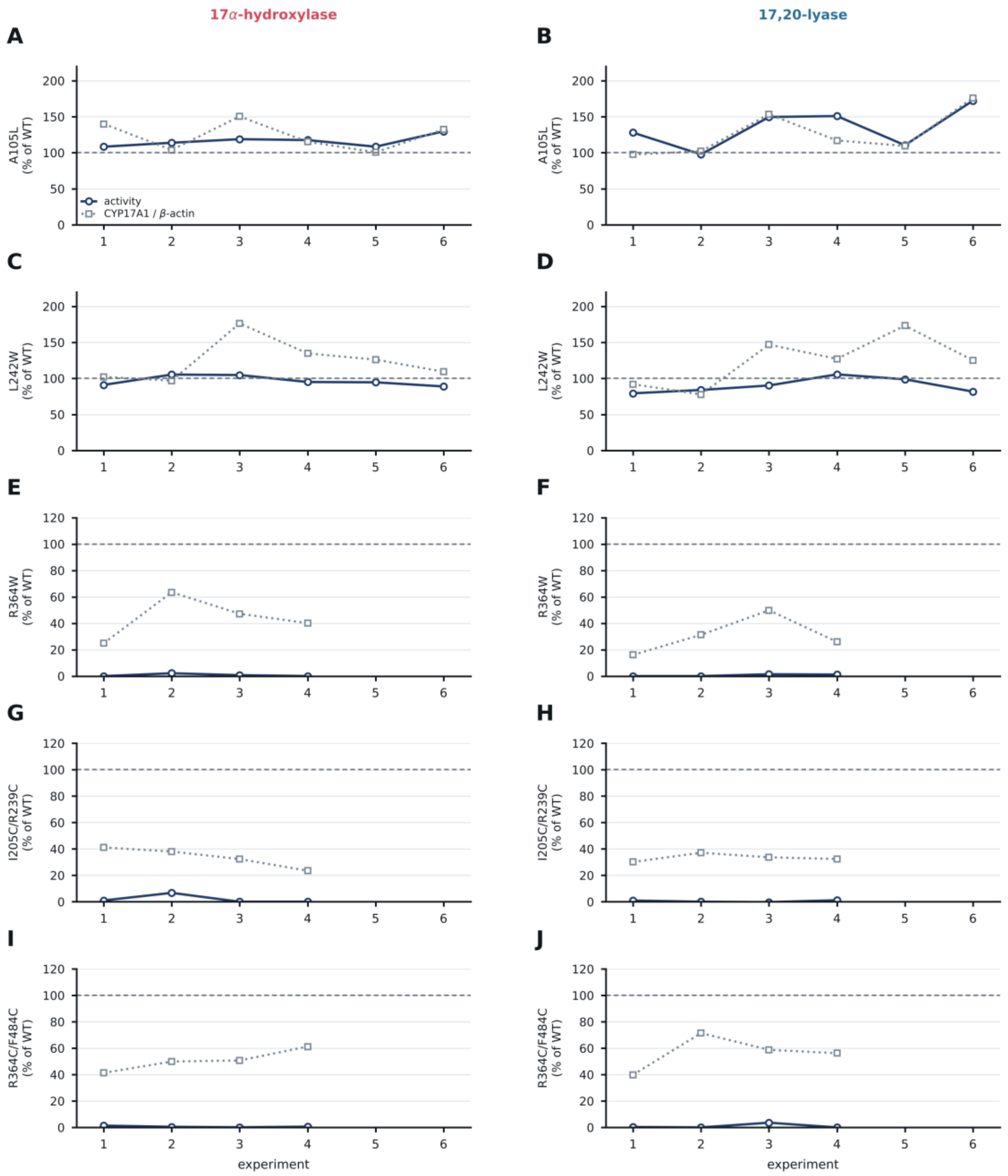


**Supplementary Figure S3.** Measured activity and CYP17A1 abundance, experiment by experiment. (A, C, E, G, I) 17α-Hydroxylase and (B, D, F, H, J) 17,20-lyase. In each panel the solid trace is the measured activity and the dotted trace the immunoblot CYP17A1/β-actin ratio of the matching lysate, both as a percentage of wild type. Note the change of vertical scale between the variants that retain activity and those that do not. Source data: Supplementary Table S9.

### Supplementary Tables

All Supplementary Tables are provided as sheets of Supplementary_Data.xlsx; sheet S0_contents lists them with the figure each supports.

Table S1. Per-cluster tunnel geometry and portal assignment (sheet S1_cluster_statistics).

Table S2. Portal-level geometry, openness and surface-forming elements (sheet S2_portal_summary).

Table S3. Median tunnel-radius profiles along the normalised path coordinate (sheet S3_radius_profiles).

Table S4. Tunnel-lining occupancy of every residue, by portal (sheet S4_lining_occupancy).

Table S5. Bottleneck-residue recurrence across the 99 tunnels (sheet S5_bottleneck_recurrence).

Table S6. Composition of the core lining of each portal (sheet S6_core_lining).

Table S7. Structural, conservation and clinical annotation of the positions discussed (sheet S7_residue_annotation).

Table S8. Design of the five portal variants (sheet S8_variant_design).

Table S9. Measured activity, immunoblot ratio and expression-corrected activity for every experiment (sheet S9_activity_per_experiment).

Table S10. Variant-level means, SD, 95% confidence intervals, CV and one-sample t-tests (sheet S10_variant_summary).

Table S11. Regression of measured activity on immunoblot abundance (sheet S11_normalisation).

Table S12. Between-experiment coefficients of variation for the measured and expression-corrected analyses (sheet S12_precision).

### Supplementary references


1. DeVore, N.M. and E.E. Scott, *Structures of cytochrome P450 17A1 with prostate cancer drugs abiraterone and TOK-001.* Nature, 2012. **482**(7383): p. 116–9.
2. Petrunak, E.M., et al., *Structures of human steroidogenic cytochrome P450 17A1 with substrates.* J Biol Chem, 2014. **289**(47): p. 32952–64.
3. Petrunak, E.M., et al., *Structural and Functional Evaluation of Clinically Relevant Inhibitors of Steroidogenic Cytochrome P450 17A1.* Drug Metab Dispos, 2017. **45**(6): p. 635–645.
4. Srejber, M., et al., *Membrane-attached mammalian cytochromes P450: An overview of the membrane's effects on structure, drug binding, and interactions with redox partners.* J Inorg Biochem, 2018. **183**: p. 117–136.

5. Fernandez-Cancio, M., et al., *Mechanism of the Dual Activities of Human CYP17A1 and Binding to Anti-Prostate Cancer Drug Abiraterone Revealed by a Novel V366M Mutation Causing 17,20 Lyase Deficiency.* Pharmaceuticals (Basel), 2018. **11**(2).
6. Pettersen, E.F., et al., *UCSF Chimera--a visualization system for exploratory research and analysis.* J Comput Chem, 2004. **25**(13): p. 1605–12.
7. Schleinkofer, K., et al., *Do mammalian cytochrome P450s show multiple ligand access pathways and ligand channelling?* EMBO Rep, 2005. **6**(6): p. 584–9.
8. Cojocaru, V., et al., *Structure and dynamics of the membrane-bound cytochrome P450 2C9.* PLoS Comput Biol, 2011. **7**(8): p. e1002152.
9. Petrek, M., et al., *CAVER: a new tool to explore routes from protein clefts, pockets and cavities.* BMC Bioinformatics, 2006. **7**: p. 316.
10. Yakubu, J., et al., *Nanoparticles with curcumin and piperine modulate steroid biosynthesis in prostate cancer.* Sci Rep, 2025. **15**(1): p. 13613.
11. Gotoh, O., *Substrate recognition sites in cytochrome P450 family 2 (CYP2) proteins inferred from comparative analyses of amino acid and coding nucleotide sequences.* J Biol Chem, 1992. **267**(1): p. 83–90.
